## Supplemental Figs S1-S4 for "The CAR mRNA-Interaction Surface is a Zipper Extension of the Ribosome A Site"

### 1. Supplementary Materials

#### *1.1 Supplementary Figures*

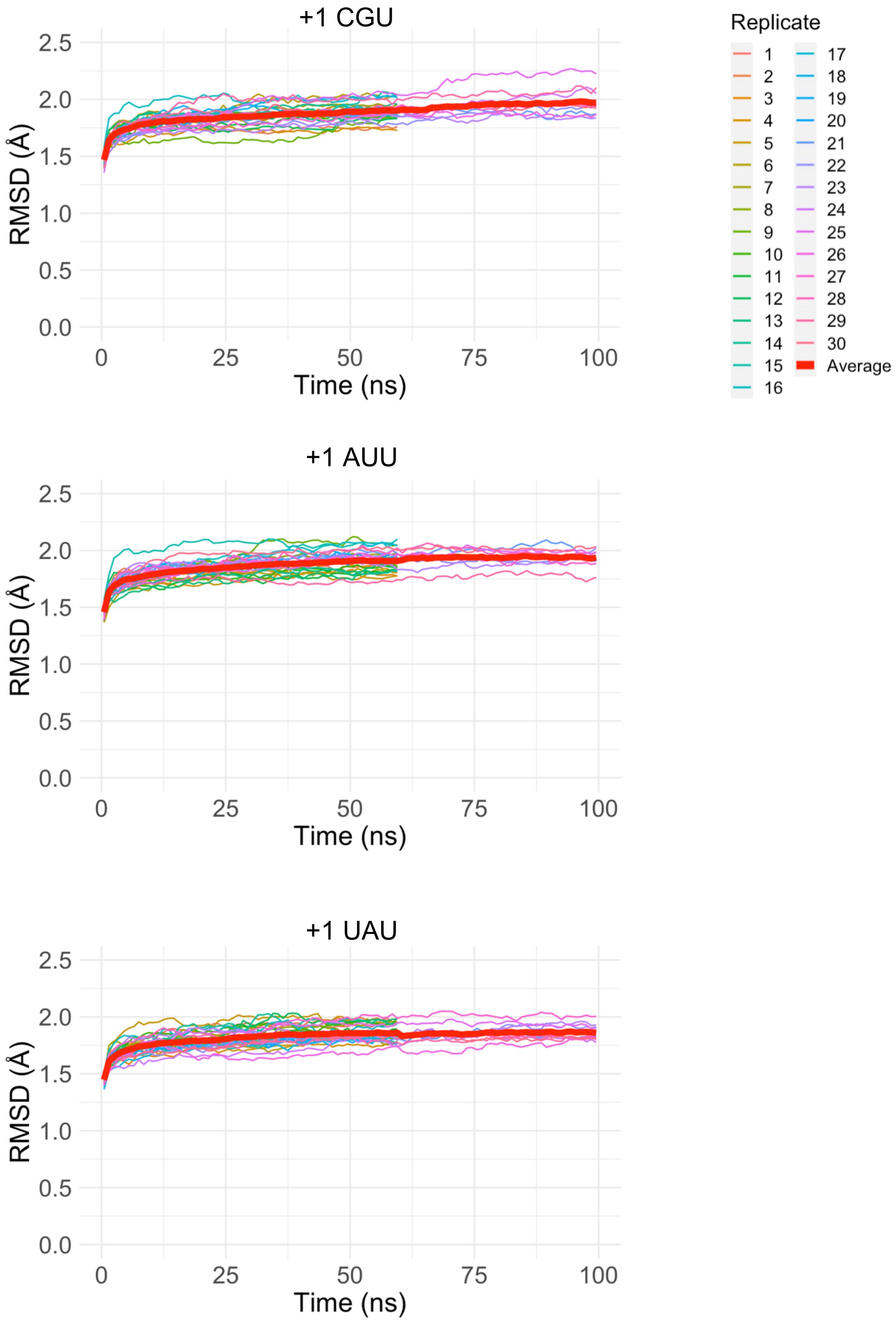


**Figure S1. RMSD profiles for trajectories.** Examples of RMSD profiles which were computed using as reference the structure at the end of equilibration. Subsystems with different +1 codons were analyzed using 60- and 100-ns trajectories. The trajectories stabilized by 20 ns and the first 20 ns of trajectories were not used for further analysis.

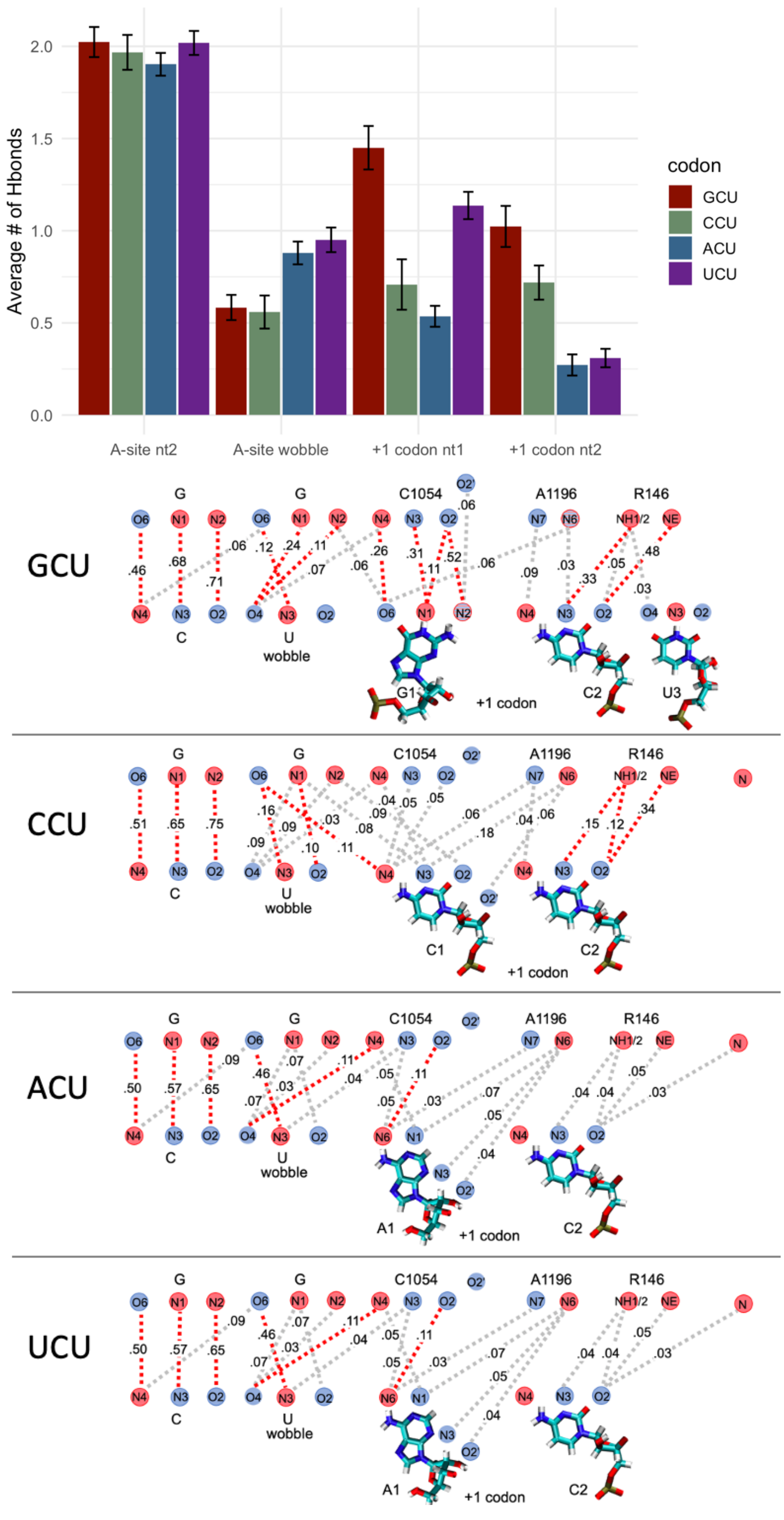


**Figure S2. H-bonding of CAR with +1 NCU codons.** +1 GCU codons have higher H-bonding with CAR compared with +1 CCU, +1 ACU and +1 UCU. Unlike the other +1 codons, H-bonding of the first nucleotide of +1 UCU is split between C1054 and A1196.

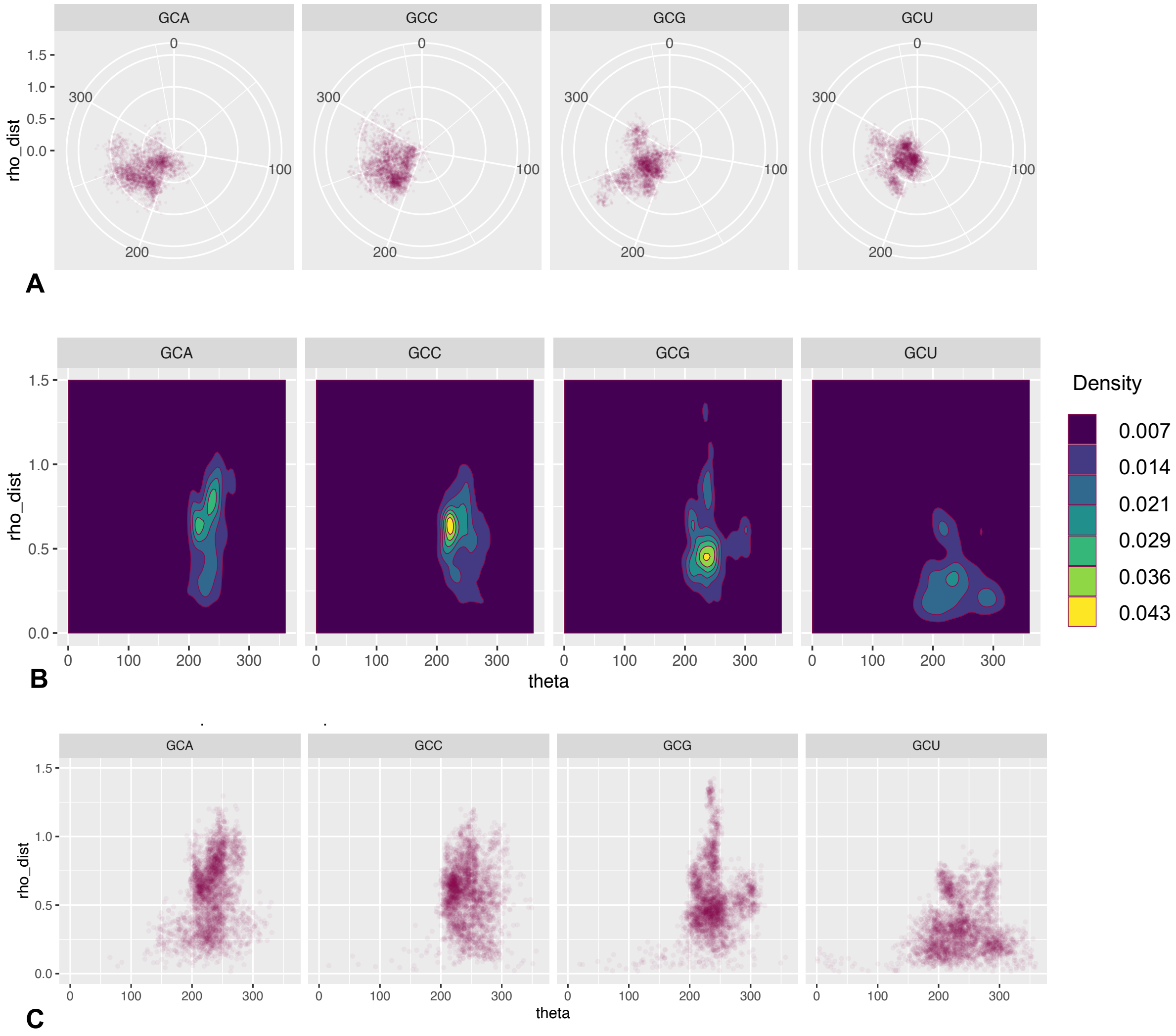


**Figure S3. Influences of nucleotide 3 of the +1 codon.** The stacking positions of G1 and C2 are compared in +1 GCA, GCC, GCG and GCU codons. The data in Figure 5D are illustrated here using scatterplot (A and C) and heat map (B) representations for the distributions of rho (Å) and theta (degrees; see Figures 5D) in 3200 frames (1600 ns) of +1 GCA, GCC, GCG and GCU trajectories. Converting the polar representation (A) to Cartesian coordinates (B and C) shows that the distribution of rho values is lower for +1 GCU codons consistent with the better centered stacking of G1 and C2.

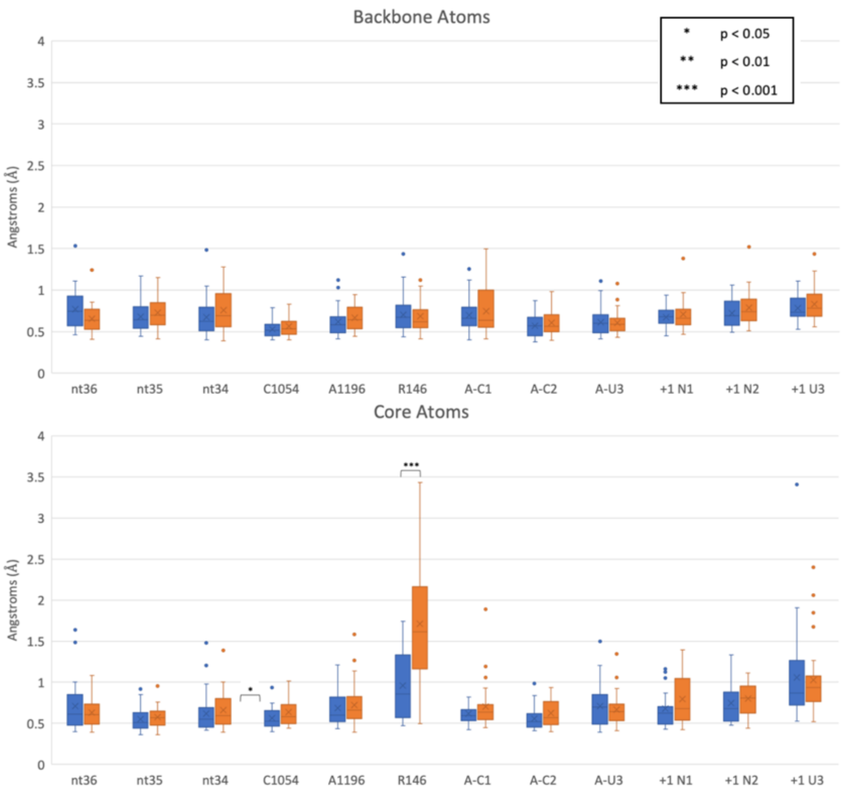
Backbone Atoms
Core Atoms
Angstroms (Å)
* p < 0.05
** p < 0.01
*** p < 0.001
nt36
nt35
nt34
C1054
A1196
R146
A-C1
A-C2
A-U3
+1 N1
+1 N2
+1 U3
0
0.5
1
1.5
2
2.5
3
3.5
4

**A**

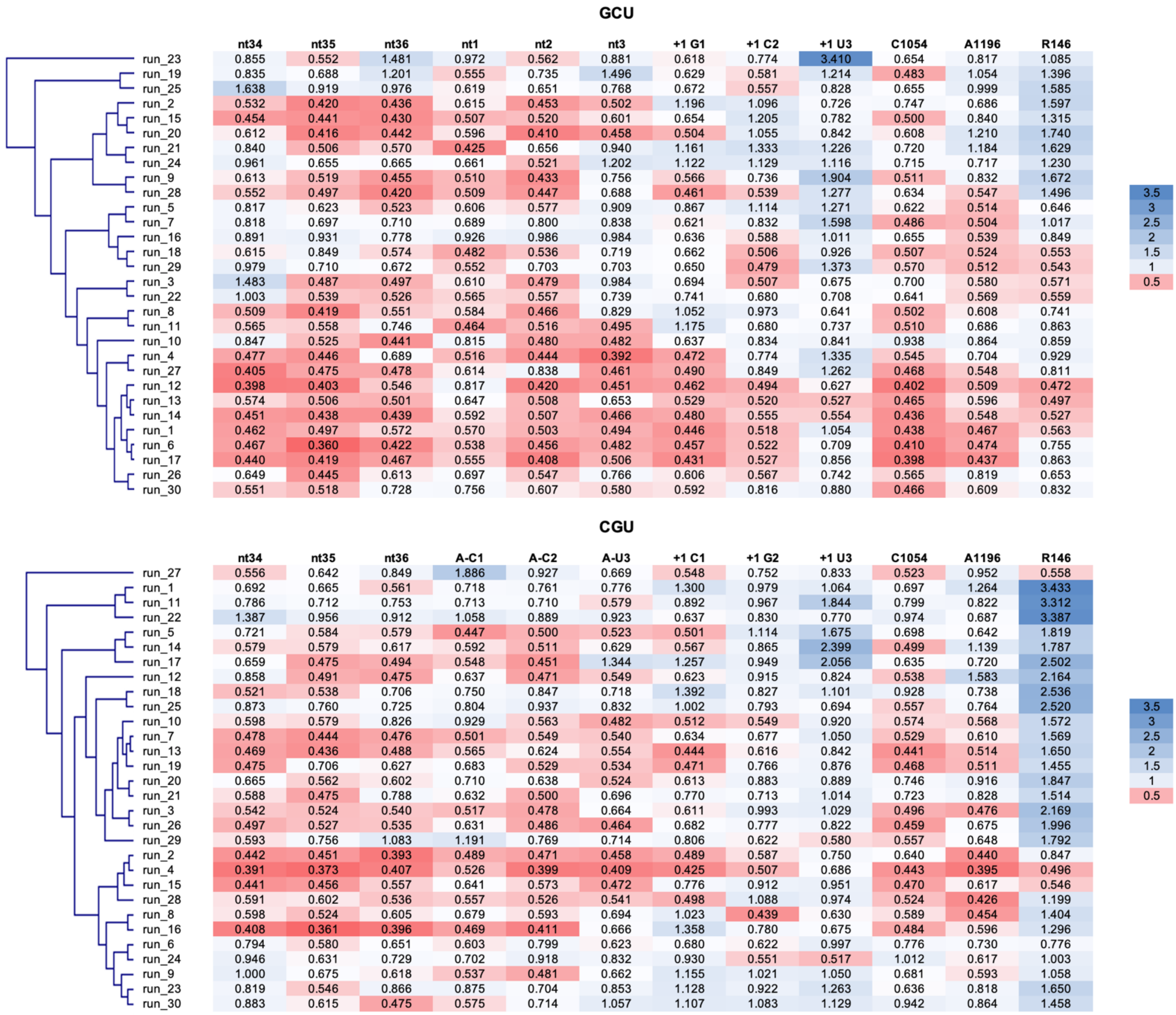


**GCU**

| | nt34 | nt35 | nt36 | nt1 | nt2 | nt3 | +1 G1 | +1 C2 | +1 U3 | C1054 | A1196 | R146 |
|---|---|---|---|---|---|---|---|---|---|---|---|---|
| run_23 | 0.855 | 0.552 | 1.481 | 0.972 | 0.562 | 0.881 | 0.618 | 0.774 | 3.410 | 0.654 | 0.817 | 1.085 |
| run_19 | 0.835 | 0.688 | 1.201 | 0.555 | 0.735 | 1.496 | 0.629 | 0.581 | 1.214 | 0.483 | 1.054 | 1.396 |
| run_25 | 1.638 | 0.919 | 0.976 | 0.619 | 0.651 | 0.768 | 0.672 | 0.557 | 0.828 | 0.655 | 0.999 | 1.585 |
| run_2 | 0.532 | 0.420 | 0.436 | 0.615 | 0.453 | 0.502 | 1.196 | 1.096 | 0.726 | 0.747 | 0.686 | 1.597 |
| run_15 | 0.454 | 0.441 | 0.430 | 0.507 | 0.520 | 0.601 | 0.654 | 1.205 | 0.782 | 0.500 | 0.840 | 1.315 |
| run_20 | 0.612 | 0.416 | 0.442 | 0.596 | 0.410 | 0.458 | 0.504 | 1.055 | 0.842 | 0.608 | 1.210 | 1.740 |
| run_21 | 0.840 | 0.506 | 0.570 | 0.425 | 0.656 | 0.940 | 1.161 | 1.333 | 1.226 | 0.720 | 1.184 | 1.629 |
| run_24 | 0.961 | 0.655 | 0.665 | 0.661 | 0.521 | 1.202 | 1.122 | 1.129 | 1.116 | 0.715 | 0.717 | 1.230 |
| run_9 | 0.613 | 0.519 | 0.455 | 0.510 | 0.433 | 0.756 | 0.566 | 0.736 | 1.904 | 0.511 | 0.832 | 1.672 |
| run_28 | 0.552 | 0.497 | 0.420 | 0.509 | 0.447 | 0.688 | 0.461 | 0.539 | 1.277 | 0.634 | 0.547 | 1.496 |
| run_5 | 0.817 | 0.623 | 0.523 | 0.606 | 0.577 | 0.909 | 0.867 | 1.114 | 1.271 | 0.622 | 0.514 | 0.646 |
| run_7 | 0.818 | 0.697 | 0.710 | 0.689 | 0.800 | 0.838 | 0.621 | 0.832 | 1.598 | 0.486 | 0.504 | 1.017 |
| run_16 | 0.891 | 0.931 | 0.778 | 0.926 | 0.986 | 0.984 | 0.636 | 0.588 | 1.011 | 0.655 | 0.539 | 0.849 |
| run_18 | 0.615 | 0.849 | 0.574 | 0.482 | 0.536 | 0.719 | 0.662 | 0.506 | 0.926 | 0.507 | 0.524 | 0.553 |
| run_29 | 0.979 | 0.710 | 0.672 | 0.552 | 0.703 | 0.703 | 0.650 | 0.479 | 1.373 | 0.570 | 0.512 | 0.543 |
| run_3 | 1.483 | 0.487 | 0.497 | 0.610 | 0.479 | 0.984 | 0.694 | 0.507 | 0.675 | 0.700 | 0.580 | 0.571 |
| run_22 | 1.003 | 0.539 | 0.526 | 0.565 | 0.557 | 0.739 | 0.741 | 0.680 | 0.708 | 0.641 | 0.569 | 0.559 |
| run_8 | 0.509 | 0.419 | 0.551 | 0.584 | 0.466 | 0.829 | 1.052 | 0.973 | 0.641 | 0.502 | 0.608 | 0.741 |
| run_11 | 0.565 | 0.558 | 0.746 | 0.464 | 0.516 | 0.495 | 1.175 | 0.680 | 0.737 | 0.510 | 0.686 | 0.863 |
| run_10 | 0.847 | 0.525 | 0.441 | 0.815 | 0.480 | 0.482 | 0.637 | 0.834 | 0.841 | 0.938 | 0.864 | 0.859 |
| run_4 | 0.477 | 0.446 | 0.689 | 0.516 | 0.444 | 0.392 | 0.472 | 0.774 | 1.335 | 0.545 | 0.704 | 0.929 |
| run_27 | 0.405 | 0.475 | 0.478 | 0.614 | 0.838 | 0.461 | 0.490 | 0.849 | 1.262 | 0.468 | 0.548 | 0.811 |
| run_12 | 0.398 | 0.403 | 0.546 | 0.817 | 0.420 | 0.451 | 0.462 | 0.494 | 0.627 | 0.402 | 0.509 | 0.472 |
| run_13 | 0.574 | 0.506 | 0.501 | 0.647 | 0.508 | 0.653 | 0.529 | 0.520 | 0.527 | 0.465 | 0.596 | 0.497 |
| run_14 | 0.451 | 0.438 | 0.439 | 0.592 | 0.507 | 0.466 | 0.480 | 0.555 | 0.554 | 0.436 | 0.548 | 0.527 |
| run_1 | 0.462 | 0.497 | 0.572 | 0.570 | 0.503 | 0.494 | 0.446 | 0.518 | 1.054 | 0.438 | 0.467 | 0.563 |
| run_6 | 0.467 | 0.360 | 0.422 | 0.538 | 0.456 | 0.482 | 0.457 | 0.522 | 0.709 | 0.410 | 0.474 | 0.755 |
| run_17 | 0.440 | 0.419 | 0.467 | 0.555 | 0.408 | 0.506 | 0.431 | 0.527 | 0.856 | 0.398 | 0.437 | 0.863 |
| run_26 | 0.649 | 0.445 | 0.613 | 0.697 | 0.547 | 0.766 | 0.606 | 0.567 | 0.742 | 0.565 | 0.819 | 0.653 |
| run_30 | 0.551 | 0.518 | 0.728 | 0.756 | 0.607 | 0.580 | 0.592 | 0.816 | 0.880 | 0.466 | 0.609 | 0.832 |

**CGU**

| | nt34 | nt35 | nt36 | A-C1 | A-C2 | A-U3 | +1 C1 | +1 G2 | +1 U3 | C1054 | A1196 | R146 |
|---|---|---|---|---|---|---|---|---|---|---|---|---|
| run_27 | 0.556 | 0.642 | 0.849 | 1.886 | 0.927 | 0.669 | 0.548 | 0.752 | 0.833 | 0.523 | 0.952 | 0.558 |
| run_1 | 0.692 | 0.665 | 0.561 | 0.718 | 0.761 | 0.776 | 1.300 | 0.979 | 1.064 | 0.697 | 1.264 | 3.433 |
| run_11 | 0.786 | 0.712 | 0.753 | 0.713 | 0.710 | 0.579 | 0.892 | 0.967 | 1.844 | 0.799 | 0.822 | 3.312 |
| run_22 | 1.387 | 0.956 | 0.912 | 1.058 | 0.889 | 0.923 | 0.637 | 0.830 | 0.770 | 0.974 | 0.687 | 3.387 |
| run_5 | 0.721 | 0.584 | 0.579 | 0.447 | 0.500 | 0.523 | 0.501 | 1.114 | 1.675 | 0.698 | 0.642 | 1.819 |
| run_14 | 0.579 | 0.579 | 0.617 | 0.592 | 0.511 | 0.629 | 0.567 | 0.865 | 2.399 | 0.499 | 1.139 | 1.787 |
| run_17 | 0.659 | 0.475 | 0.494 | 0.548 | 0.451 | 1.344 | 1.257 | 0.949 | 2.056 | 0.635 | 0.720 | 2.502 |
| run_12 | 0.858 | 0.491 | 0.475 | 0.637 | 0.471 | 0.549 | 0.623 | 0.915 | 0.824 | 0.538 | 1.583 | 2.164 |
| run_18 | 0.521 | 0.538 | 0.706 | 0.750 | 0.847 | 0.718 | 1.392 | 0.827 | 1.101 | 0.928 | 0.738 | 2.536 |
| run_25 | 0.873 | 0.760 | 0.725 | 0.804 | 0.937 | 0.832 | 1.002 | 0.793 | 0.694 | 0.557 | 0.764 | 2.520 |
| run_10 | 0.598 | 0.579 | 0.826 | 0.929 | 0.563 | 0.482 | 0.512 | 0.549 | 0.920 | 0.574 | 0.568 | 1.572 |
| run_7 | 0.478 | 0.444 | 0.476 | 0.501 | 0.549 | 0.540 | 0.634 | 0.677 | 1.050 | 0.529 | 0.610 | 1.569 |
| run_13 | 0.469 | 0.436 | 0.488 | 0.565 | 0.624 | 0.554 | 0.444 | 0.616 | 0.842 | 0.441 | 0.514 | 1.650 |
| run_19 | 0.475 | 0.706 | 0.627 | 0.683 | 0.529 | 0.534 | 0.471 | 0.766 | 0.876 | 0.468 | 0.511 | 1.455 |
| run_20 | 0.665 | 0.562 | 0.602 | 0.710 | 0.638 | 0.524 | 0.613 | 0.883 | 0.889 | 0.746 | 0.916 | 1.847 |
| run_21 | 0.588 | 0.475 | 0.788 | 0.632 | 0.500 | 0.696 | 0.770 | 0.713 | 1.014 | 0.723 | 0.828 | 1.514 |
| run_3 | 0.542 | 0.524 | 0.540 | 0.517 | 0.478 | 0.664 | 0.611 | 0.993 | 1.029 | 0.496 | 0.476 | 2.169 |
| run_26 | 0.497 | 0.527 | 0.535 | 0.631 | 0.486 | 0.464 | 0.682 | 0.777 | 0.822 | 0.459 | 0.675 | 1.996 |
| run_29 | 0.593 | 0.756 | 1.083 | 1.191 | 0.769 | 0.714 | 0.806 | 0.622 | 0.580 | 0.557 | 0.648 | 1.792 |
| run_2 | 0.442 | 0.451 | 0.393 | 0.489 | 0.471 | 0.458 | 0.489 | 0.587 | 0.750 | 0.640 | 0.440 | 0.847 |
| run_4 | 0.391 | 0.373 | 0.407 | 0.526 | 0.399 | 0.409 | 0.425 | 0.507 | 0.686 | 0.443 | 0.395 | 0.496 |
| run_15 | 0.441 | 0.456 | 0.557 | 0.641 | 0.573 | 0.472 | 0.776 | 0.912 | 0.951 | 0.470 | 0.617 | 0.546 |
| run_28 | 0.591 | 0.602 | 0.536 | 0.557 | 0.526 | 0.541 | 0.498 | 1.088 | 0.974 | 0.524 | 0.426 | 1.199 |
| run_8 | 0.598 | 0.524 | 0.605 | 0.679 | 0.593 | 0.694 | 1.023 | 0.439 | 0.630 | 0.589 | 0.454 | 1.404 |
| run_16 | 0.408 | 0.361 | 0.396 | 0.469 | 0.411 | 0.666 | 1.358 | 0.780 | 0.675 | 0.484 | 0.596 | 1.296 |
| run_6 | 0.794 | 0.580 | 0.651 | 0.603 | 0.799 | 0.623 | 0.680 | 0.622 | 0.997 | 0.776 | 0.730 | 0.776 |
| run_24 | 0.946 | 0.631 | 0.729 | 0.702 | 0.918 | 0.832 | 0.930 | 0.551 | 0.517 | 1.012 | 0.617 | 1.003 |
| run_9 | 1.000 | 0.675 | 0.618 | 0.537 | 0.481 | 0.662 | 1.155 | 1.021 | 1.050 | 0.681 | 0.593 | 1.058 |
| run_23 | 0.819 | 0.546 | 0.866 | 0.875 | 0.704 | 0.853 | 1.128 | 0.922 | 1.263 | 0.636 | 0.818 | 1.650 |
| run_30 | 0.883 | 0.615 | 0.475 | 0.575 | 0.714 | 1.057 | 1.107 | 1.083 | 1.129 | 0.942 | 0.864 | 1.458 |

**B**

**Figure S4. RMSF comparison of +1 GCU and +1 CGU trajectories.** (A) R146 in +1 CGU trajectories has elevated RMSF (blue) compared to +1 GCU (orange; $p < 0.001$ ***). RMSF was measured using "core" base heavy atoms of nucleotide bases (C2, C4, C5, C6, N1, N3 of C and U; C2, C4, C5, C6, C8, N1, N3, N7, and G) and the guanidinium group (CZ, NE, NH1, NH2). (B) RMSF measurements for each MD experiment (row).
